# ^1^H MR Spectroscopy of the Motor Cortex Immediately following Transcranial Direct Current Stimulation at 7 Tesla

**DOI:** 10.1101/322941

**Authors:** Kayla Ryan, Krzysztof Wawrzyn, Joseph Gati, Blaine A. Chronik, Dickson Wong, Neil Duggal, Robert Bartha

**Author notes:** Corresponding Author: Robert Bartha, PhD Centre for Functional and Metabolic Mapping (CFMM), Robarts Research Institute, The University of Western Ontario, 1151 Richmond Street, London, Ontario, Canada N6A 3K7.

## Abstract

Transcranial direct current stimulation (tDCS) is a form of non-invasive brain stimulation that may modulate cortical excitability, metabolite concentration, and human behaviour. The supplementary motor area (SMA) has been largely ignored as a potential target for tDCS neurorehabilitation but is an important region in motor compensation after brain injury with strong efferent connections to the primary motor cortex (M1). The objective of this work was to measure tissue metabolite changes in the human motor cortex immediately following tDCS. We hypothesized that bihemispheric tDCS would change levels of metabolites involved in neuromodulation including *N*-acetylaspartate, glutamate, and creatine. In this single-blind, randomized, cross-over study, fifteen healthy adults aged 21-60 participated in two 7T MRI sessions, to identify changes in metabolite concentrations by magnetic resonance spectroscopy. Immediately after 20 minutes of tDCS, there were no significant changes in metabolite levels or metabolite ratios comparing tDCS to sham. However there was a trend toward increased NAA/tCr concentration (*p*=0.08) in M1 under the stimulating cathode. There was a strong, positive correlation between the change in the absolute concentration of NAA and the change in the absolute concentration of tCr (*p*<0.001) suggesting an effect of tDCS. Both NAA and creatine are important markers of neurometabolism. Our findings provide novel insight into the modulation of neural metabolites in the motor cortex *immediately following* application of bihemispheric tDCS.

## Introduction

Transcranial direct current stimulation (tDCS) is a form of non-invasive brain stimulation that has shown promise in modulating cortical excitability and behaviour in humans (1–4). However, many facets of its use remain controversial. For example, the optimization of stimulation parameters (current level, duration, electrode montage, etc.), characterization of individual response variability, and the physiological mechanism of action are still under active investigation.

Several potential mechanisms of action have been proposed based on pharmacological, behavioural, and imaging studies (1–3, 5, 6). At current levels of ~1 mA, tDCS is thought to induce alterations of the membrane potential, with anodal tDCS making it more likely, and cathodal tDCS making it less likely, for an action potential to fire (7, 8). However, the level of current may also impact the neuronal response to stimulation. At 2 mA, cathodal stimulation has been shown to have an excitatory influence on membrane potential. This has been thought to occur due to an increased release of Ca^2+^ at the higher current (9). Furthermore, higher current penetrates deeper into cortical tissues, potentially causing dendritic depolarization at a sufficient level to modulate excitation of adjacent neuronal structures (9). Magnetic resonance spectroscopy (MRS) provides a means to investigate the effects of tDCS on cellular metabolism and synaptic transmission as it can be used to non-invasively quantify cerebral metabolites *in vivo*, including glutamate (Glu) and gamma-aminobutyric acid (GABA). Previous MRS studies have shown changes in excitatory and inhibitory neurotransmitter levels, minutes after tDCS, with current levels ranging from 1-2 mA (6, 7, 10–14). Other studies have suggested that creatine may have an important role in bioenergetics and neuromodulation (15–17). For example, Rae *et al.* found an increase in adenosine-triphosphate (ATP) synthesis, with a decrease in the concentration of phosphocreatine in the left temporo-frontal region following anodal tDCS to the left dorsolateral prefrontal cortex (17). In another study, 2mA of anodal tDCS to the right parietal cortex caused an increase in both Glx and total *N*-aceytl-aspartate (NAA + NAAG) relative to sham, measured from the parietal cortex (10), while a study by Stagg and colleagues found that 1 mA of cathodal stimulation to left M1 decreased Glx under the electrode(7). Other studies have found no effect. For example, Kim *et. al.* found no changes after 1.5 mA of cathodal tDCS to left M1 in any metabolite measured under the stimulating electrode (18). Similarly, using 1mA of current in an M1-M1 bihemispheric montage, Tremblay *et. al.* found no significant changes in any metabolite in left M1 (19). These conflicting results are difficult to interpret, and leads to uncertainty with regards to the implementation of an optimum stimulation paradigm.

The application of tDCS to improve motor performance and recovery in neurological disorders requires optimization of stimulation parameters. Bihemispheric tDCS can enhance both behaviour and physiological responses in healthy and neurologically injured individuals (20–22). The supplementary motor area (SMA) has proven to be an important area of the brain during the execution of bimanual hand movements (23), and plays a compensatory role during the recovery of both stroke and spinal cord injury (24–26). With its strongest efferent projections to M1 and the corticospinal tract, SMA is a unique target for tDCS (27). Support for this notion comes from a recent study that showed enhanced motor performance by targeting the left SMA with 0.4 mA of anodal tDCS for 90 min over three days (28). By targeting *both SMA and M1* with 2mA of tDCS, it may be possible to induce additive effects on M1 excitability via interhemispheric connections, which are thought to be more focal than those associated with M1-supraortibal stimulation (19, 20).

The purpose of the current study was to demonstrate the feasibility of concurrent tDCS and 7T MRI, and to determine whether targeting both SMA and M1 using a bihemispheric tDCS montage would produce immediate changes in metabolite concentrations in M1 measured using ultra high-field (7T) MRS. To our knowledge, this is the first study to examine the metabolic changes after bihemispheric tDCS, delivered in the MR environment, at an ultra-high magnetic field strength. Based on previous studies, we hypothesized that 2 mA bihemispheric tDCS would enhance synaptic and metabolic activity (10, 11, 15, 17). As such, metabolites involved in neurometabolism such as NAA, glutamate, and creatine would be altered by stimulation.

## Methods

### Participants and Study Design

15 healthy adults aged 21-60 years (mean ± standard deviation: 28 ± 10, 9 female), with no reported history of mental or neurological illness, participated in two sessions on a 7 Tesla (Siemens, Erlangen) head-only MRI scanner. All participants had ^1^H MRS in this single blind, sham controlled, cross-over design. Participants were randomized to receive tDCS stimulation or sham stimulation on their initial visit, and the contrary on their second visit, at least 7 days apart. Informed written consent was obtained for all procedures according to the Declaration of Helinski (World Medical Association, 2008) and the study was approved by the Western University Health Sciences Research Ethics Board.

### tDCS Stimulation

Using an MR-compatible DC-STIMULATOR (NeuroConn, Germany), 2 mA of current was applied to bihemispheric motor areas in the MRI scanner, for a total of 20 minutes.

Electrodes were 3×3 cm^2^, providing a total current density of 0.22 mA/cm^2^ and a total charge with respect to time of 0.33 C/cm^2^. For use inside the scanner, electrodes were fit with 5 kOhm resistors placed next to the electrode to minimize the possibility of eddy currents induced in the leads during MRS acquisition. Electrodes were positioned on each participant outside the magnet using the EEG 10-10 system, which has been shown to be a reliable localization tool (29). The cathode was placed on the left primary motor cortex (C3), anode on the right supplementary motor area (FC_2_). For stimulation, current was ramped up over 10 seconds to reach 2 mA and held constant for 20 minutes, followed by a 10 s ramp down period. During sham stimulation, current was ramped up over 10 s and then immediately turned off. As it has been shown that subjects are unable to distinguish between sham and true tDCS using this paradigm, we used this measurement as a baseline comparison (3, 30).

### Temperature Monitoring

To ensure the safety of the participants during tDCS in the MRI, temperature was monitored on all subjects throughout the duration of the scan (approximately an hour and 15 minutes). Specifically, four T1C 1.7 mm diameter fibre optic temperature sensors (Neoptix, Quebec, Canada) were located under both electrode pads and the nearest cable chokes. Temperature was monitored in real time with a calibrated Reflex signal conditioner (Neoptix, Quebec, Canada) and a custom data collection program written in LabVIEW 2010 (National Instruments).

### Magnetic Resonance Image Acquisition and Analysis

A 7 Tesla Siemens (Erlangen, Germany), head-only MRI (Magnetom) was used to acquire spectroscopy and imaging data. Data were acquired using an 8 channel transmit and 32 channel receive coil array. T_1_-weighted MP2RAGE anatomical images (TE/TR = 2.83/6000 ms and 750 μm isotropic resolution) were acquired and used for voxel positioning. These images were also used to estimate white-matter (WM), gray-matter (GM) and cerebrospinal fluid (CSF) fractions for partial volume correction when determining metabolite concentration. The MRS acquisition began immediately following the completion of the stimulation to capture alterations in metabolite concentration due to tDCS. Water suppressed (64 averages) and unsuppressed (8 averages) ^1^H MR spectra were acquired from a single voxel (1.6×2.0×1.8 cm^3^) located in the left primary motor cortex (under the cathode) (Figure 1) using the semi Localization by Adiabatic Selective Refocusing (semi-LASER) pulse sequence (31): TE/TR = 60/7500 ms, voxel size=1.6×2.0×1.8 cm^3^,total MRS acquisition time was approximately 10 minutes. A localized B_0_ and B_1_ shim were applied prior to data acquisition. The B_0_ shim was optimized using a two-echo gradient recalled echo (GRE) shimming technique (32) and the B_1_ field was optimized such that the phases of the transmit channels added constructively within the MRS voxel. Spectra were lineshape corrected using combined QUALITY deconvolution and eddy current correction (QUECC) with 400 QUALITY points (33). Simulated prior knowledge metabolite lineshapes were fitted to post-processed spectra using the fitMAN software developed in-house (Figure 2) (34). Metabolite concentrations were examined as ratios normalized to creatine and also as absolute concentrations using unsuppressed water as an internal reference standard as previously described (35). Measurement of tissue partial volume with the voxel was made using the MP2RAGE images in FMRIB Software Library (FSL) (36) to obtain the fraction of WM, GM and CSF within the voxel. In addition, relaxation rates of the metabolites were incorporated into the quantification to correct for T_1_ and T_2_ relaxation induced signal loss (37–40).

**Figure 1.**
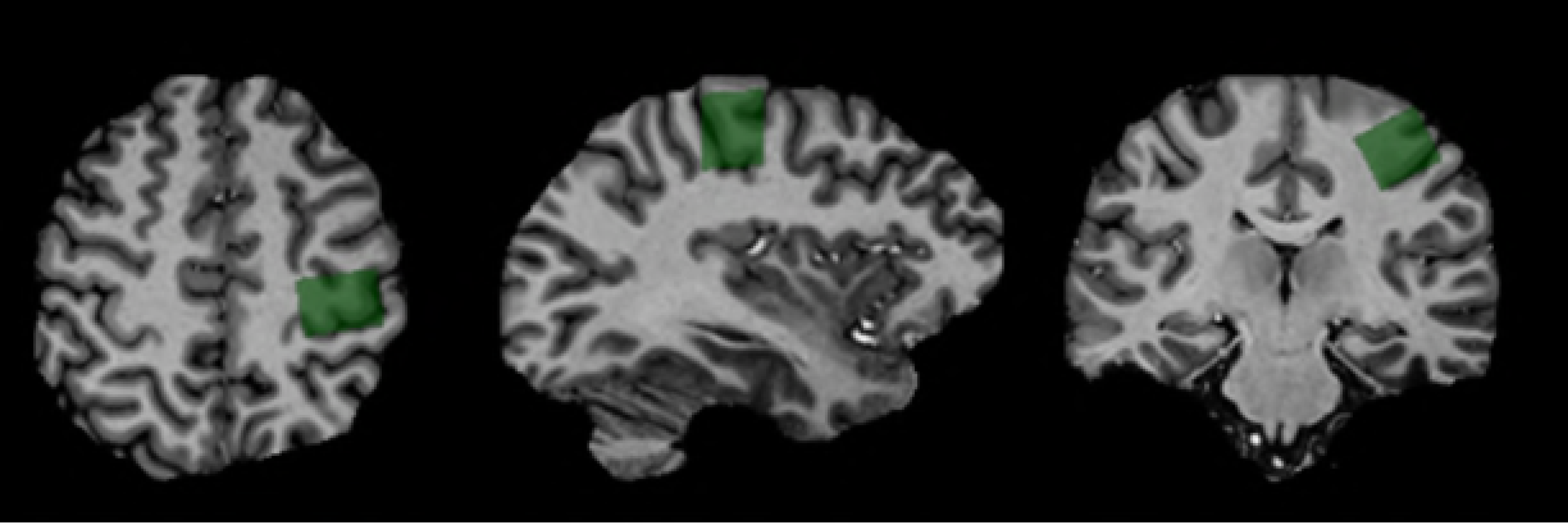
Voxel Positioning: Typical MP2RAGE anatomical images used for voxel placement were brain extracted using FSL. The voxel shown in green (2×2×2 cm^3^) was placed over the left primary motor cortex (under the cathode).

**Figure 2.**
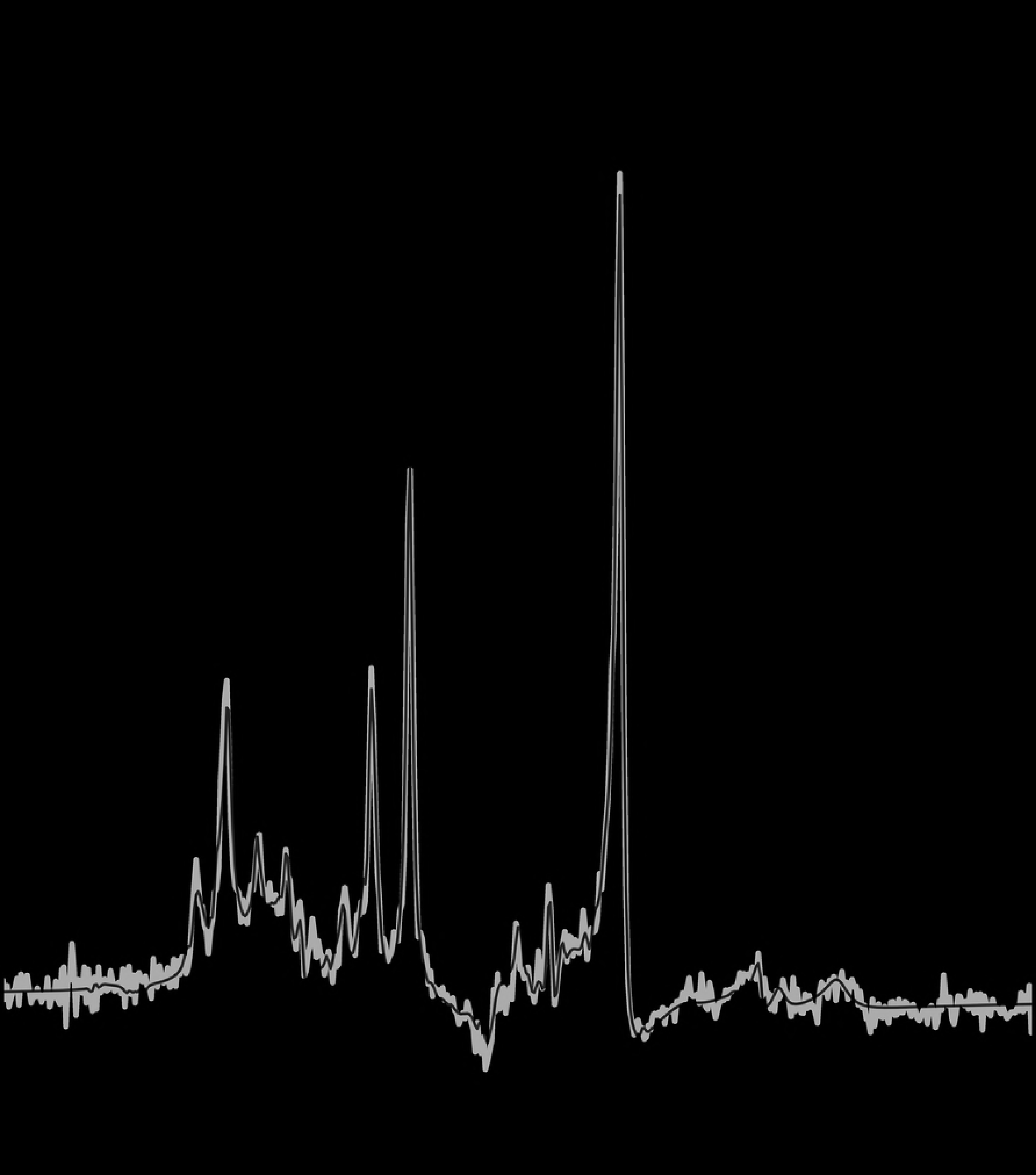
Spectrum of Left Motor Cortex: 7 Tesla semi-LASER ^1^H MRS (TE=60 ms) of the left primary motor cortex. The spectrum (grey) is overlaid on the fitted result (black) with the residual shown above (black). Select metabolite peaks are identified.

Metabolites measured with a group coefficient of variation of less than <30% in the sham condition were included in statistical analyses. To identify differences in metabolite levels and metabolite ratios between sham and tDCS conditions, repeated measures MANOVA was performed in SPSS (IBM SPSS Statistics Version 25). The main factors were the type of stimulation (two levels: sham and tDCS) and the metabolite (six levels: *N*-acetyl aspartate (NAA), myo-inositol (mI), creatine (Cr), choline (Cho), glutamate (Glu), glutathione (GSH) or metabolite ratio (five levels: NAA/Cr, mI/Cr, Cho/Cr, Glu/Cr, GSH/Cr) measured. In addition, differences between sham and tDCS conditions were compared separately for each metabolite and metabolite ratio using paired t-tests.

## Results

### Temperature Monitoring

The tDCS was safely and successfully applied in the 7T MRI environment in all subjects. The average temperature change in all four probes was 4.3 ± 0.2 °C throughout the duration of the experiment. This temperature increase was largely due to warming of the bore and from the participant’s natural body heating. Once equilibrium was established, small fluctuations on the order of 1 °C were observed during periods when RF was turned on.

### Metabolite Ratio Changes

Spectral quality measures including signal to noise ration and linewidth are summarized in Table 1 for all participants. There were no age or gender related effects. When examining the metabolite ratios the repeated measures MANOVA indicated a trend for the effect of stimulation (F_(1,14)_ =3.52, *p*=0.08). In addition, there was a significant main effect of metabolite (F_(12,3)_ =343.35, *p*<0.001). There was no main interaction effect (F_(12,3)_=1.25, *p*=0.33) (Table 2). *Post-hoc* t-tests (uncorrected for multiple comparisons) were performed to confirm alterations in metabolite ratios. Our results showed a trend toward a 4% increase in the NAA/tCr ratio between sham and tDCS conditions (*p*=0.08, Cohen’s d =0.52) and no significant changes in any other metabolite ratios (Figure 3, Table 2).

**Table 1.** Spectral Quality and Quantification: Characterization of spectral quality and voxel tissue composition. Signal to noise ratio represents the intensity of the NAA_Ch3_ peak divided by the standard deviation of the baseline noise after Fourier transformation of the initial 0.3 seconds. The linewidth represents the full width at half maximum (FWHM) of the unsuppressed water signal. The water area represents the area of the unsuppressed water spectrum. The voxel tissue partial volume is provided for gray matter (GM), white matter (WM) and cerebral spinal fluid (CSF). Data is presented as mean ± standard error of the mean. Repeated measured t-tests were conducted on all spectral parameters; no significant changes between sham and stimulation were observed.

|  | **Sham** | **tDCS** |
| --- | --- | --- |
| Signal to noise ratio | $78.9 \pm 5.2$ | $82.3 \pm 4.6$ |
| Linewidth (Hz) | $13.5 \pm 1.2$ | $13.6 \pm 1.3$ |
| Water Area | $4.8 \pm 0.16$ | $4.9 \pm 0.02$ |
| GM Fraction | $0.37 \pm 0.01$ | $0.37 \pm 0.02$ |
| WM Fraction | $0.53 \pm 0.02$ | $0.53 \pm 0.03$ |
| CSF Fraction | $0.09 \pm 0.008$ | $0.10 \pm 0.009$ |

**Table 2.** Analysis of Metabolic Ratios: Metabolite ratios relative to total creatine. The *p*-values were calculated in post-hoc analysis of individual metabolite ratios using paired t-tests.

|  | **Sham** | **tDCS** | ***p*-value** |
| --- | --- | --- | --- |
| NAA/Cr | 1.67 ± 0.03 | 1.73 ± 0.03 | 0.08 |
| Cho/Cr | 0.68 ± 0.02 | 0.67± 0.02 | 0.78 |
| Myo/Cr | 0.72 ± 0.02 | 0.72 ± 0.02 | 0.79 |
| Glu/Cr | 0.71 ± 0.06 | 0.79 ± 0.07 | 0.21 |
| GSH/Cr | 1.28 ± 0.05 | 1.40 ± 0.06 | 0.21 |

**Figure 3.**
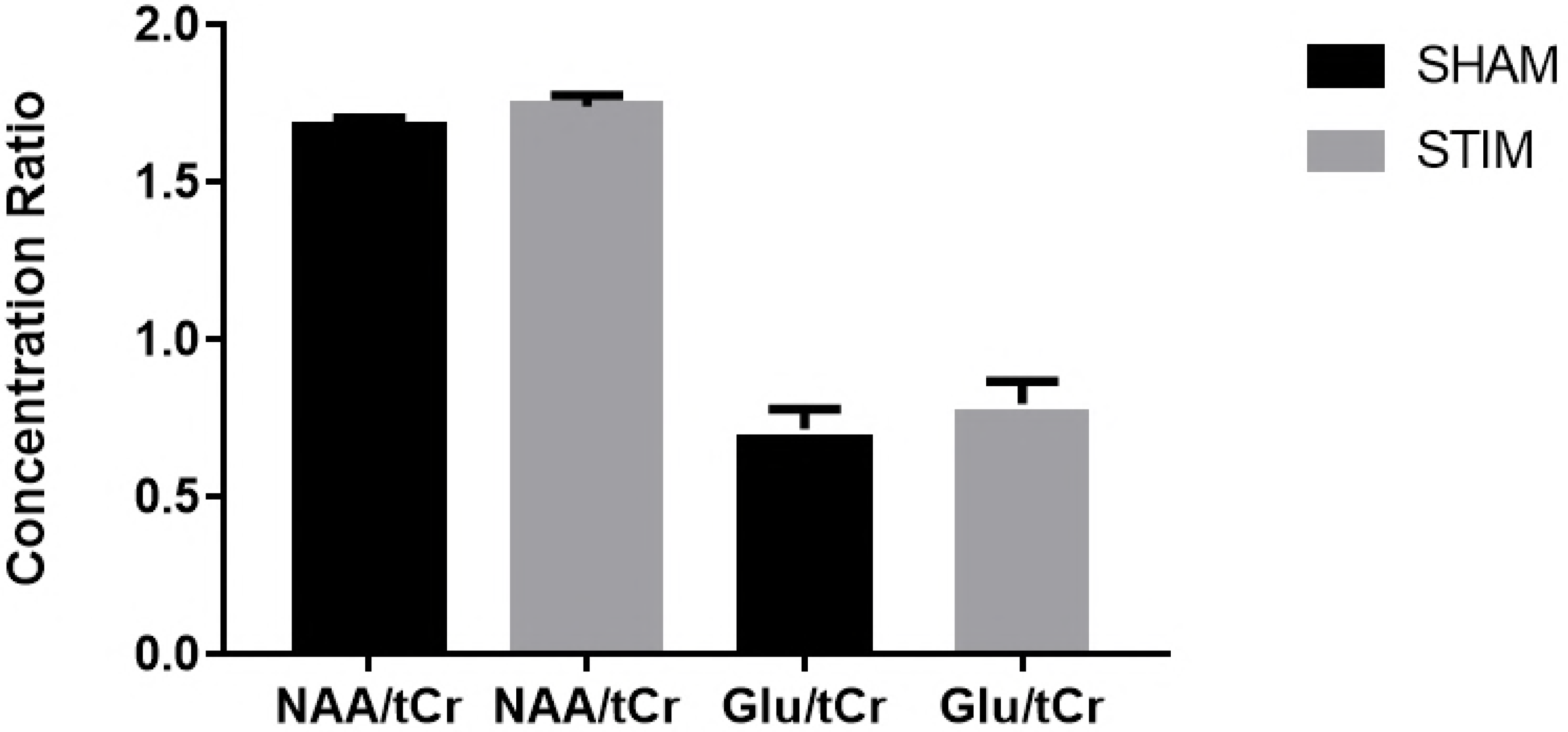
Metabolic ratios - sham vs stim: tDCS increases NAA/tCr ratio when measured immediately after a 20 minute stimulation period (* *p*=0.08). No difference was observed in Glu/Cr. Error bars indicate SEM.

### Metabolite Concentration Changes

Using repeated measures MANOVA, there was no main effect of bihemispheric M1-SMA tDCS on the absolute concentration of any metabolite (F_(1,14)_=1.55, *p*=0.23). There was also no significant interaction effect of metabolite and condition (F_(4,11)_ =1.42, *p*=0.29). Table 3 displays the absolute metabolite concentrations. *Post-hoc* comparisons using paired t-tests (uncorrected for multiple comparisons) showed a trend toward decreased tCr (*p*=0.07, Cohen’s d =0.42) and decreased mI (*p*=0.08, Cohen’s d=0.48), with no significant changes in any metabolites.

**Table 3.** Analysis of Absolute Metabolic Concentration: Absolute concentration of metabolites measured by MRS. The *p*-values were measured by post-hoc analysis of individual metabolites using paired t-tests.

| Metabolite | Sham tDCS | tDCS | *p*-value |
| --- | --- | --- | --- |
| NAA (mM) | 16.2 ± 0.65 | 15.6 ± 0.46 | 0.40 |
| Cho (mM) | 2.4 ± 0.1 | 2.3 ± 0.11 | 0.14 |
| Myo (mM) | 6.4 ± 0.28 | 5.9 ± 0.24 | 0.08 |
| tCr (mM) | 10.9 ± 0.4 | 10.1 ± 0.3 | 0.07 |
| Glu (mM) | 7.9 ± 0.71 | 8.2 ± 0.79 | 0.69 |
| GSH (mM) | 2.4 ± 0.18 | 2.4 ± 0.13 | 0.88 |

### Correlation between NAA and tCr

We observed a strong, positive correlation between the change in the absolute concentration of NAA and the change in the absolute concentration of tCr (stimulation - sham, R^2^ = 0.64, p < 0.001, Figure 4).

**Figure 4.**
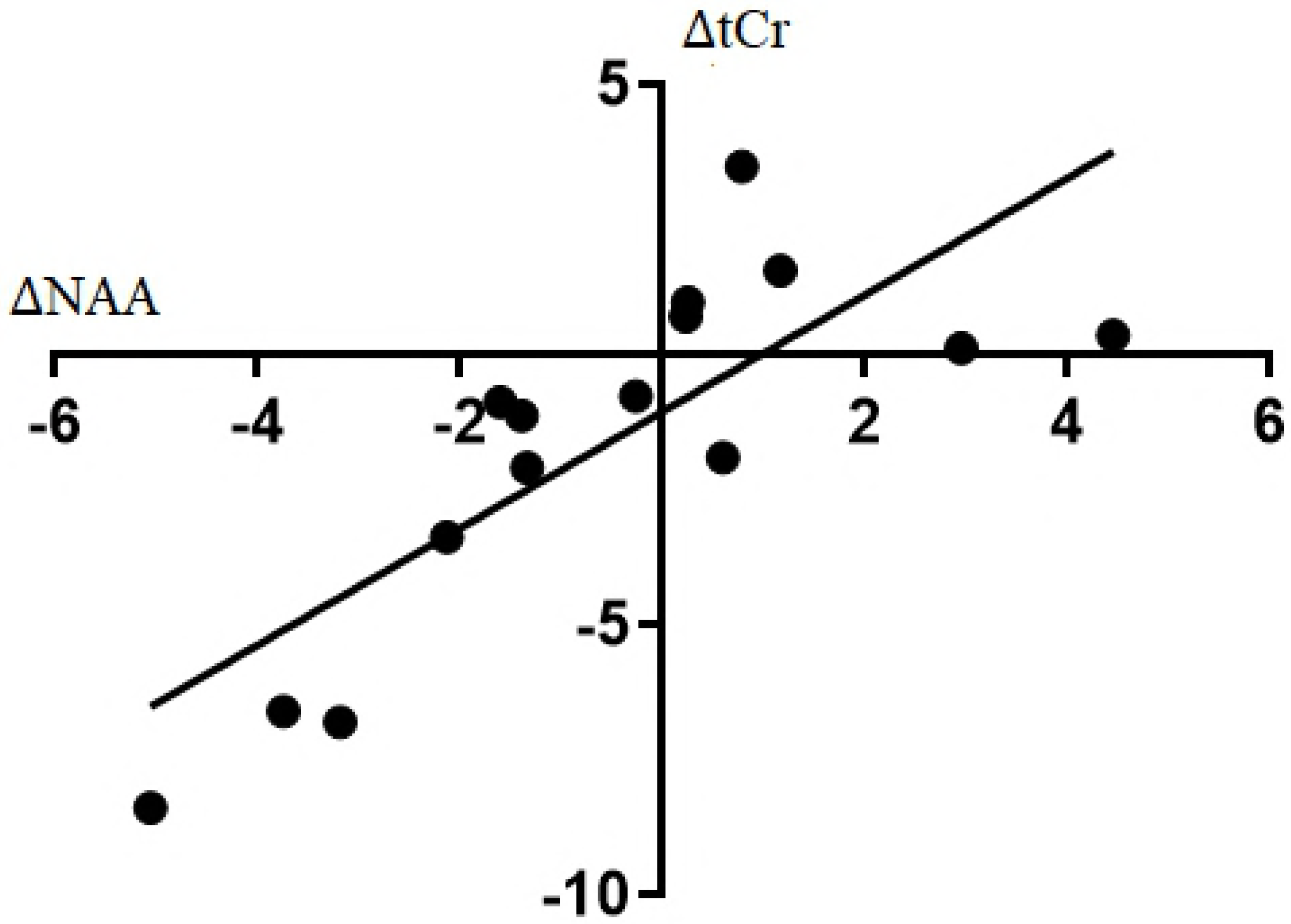
NAA and tCr Coupling: The association between the Δ in the absolute concentration of NAA and tCr (stimulation - sham). We observed a strong, positive correlation, indicating NAA and tCr both change in the same direction after stimulation (R^2^ = 0.64, p <0.001).

## Discussion

Bihemispheric tDCS was safely and successfully performed in the 7T MRI environment with minimal heating effects. However, the bihemispheric tDCS of M1-SMA produced no significant metabolite level changes in the left primary motor cortex (M1) immediately after 20 minutes of stimulation measured by 7T MR spectroscopy. *Post-hoc* analysis did show a trend toward increased NAA/tCr and decreased levels of tCr. In addition, we observed a strong association between the change in absolute concentration of NAA and the change in absolute concentration of tCr that may indicate a coupling between these metabolites following tDCS.

The trends toward lower NAA/tCr and tCr observed in the current study may be due to changes in brain activity induced by tDCS. Brain activity has been shown to decrease both NAA and tCr levels. Specifically, NAA is associated with metabolic and mitochondrial activity (16, 41). Following visual stimulation, Baslow and colleagues found that the concentration of NAA decreased by approximately 13% in the visual cortex (42). Similarly, Castellano and colleagues observed a 20% decrease in NAA after visual stimulation (43). This decrease in NAA was attributed to a lower rate of NAA synthesis compared to hydrolysis during periods of cortical activation, suggesting that the brain used NAA faster than it could be synthesized (42, 43). NAA is the precursor for the synthesis of *N*-acetylaspartylglutamate (NAAG), a modulator of glutamate and GABA neurotransmitter release. When neural activity is increased, there is an increased release of NAAG from the synapse (43, 44). It has been suggested that the reduction in NAA upon neural activation is due to increased demand for NAAG. In support of this hypothesis, both Landim *et al.* and Castellano *et al.* observed a decrease in NAA concentration with a subsequent increase in NAAG upon stimulation (43, 44).

Creatine (Cr) may also be altered as a consequence of neuronal stimulation due to its role in energy metabolism through its conversion to phosphocreatine (PCr) (15, 45). In the central nervous system, Cr and PCr are involved in maintaining the high energy levels necessary for the maintenance of membrane potentials, ion gradients, calcium homeostasis, and intracellular signalling (46). Cr has also been observed as a potential modulator of neurotransmission (15, 45). The Cr peak measured by MR spectroscopy represents intracellular contributions from both Cr and PCr (tCr). In areas of high energy demand, PCr is used to convert ADP to ATP. As such, intracellular stores of PCr will transiently decrease, consistent with the trend toward decreased tCr observed in the current study. Rango *et. al.* have also discovered a transient decrease in PCr after short bursts of visual stimulation, concluding that functional activation reduces PCr (47). Furthermore, Cr is released from the neuron in an action potential dependent manner. An increase in the resting membrane potential, induced by tDCS, may result in release of Cr from the neuron to act as a co-transmitter. Early studies on rodents indicate Cr may modulate postsynaptic neurotransmitters such as GABA, inhibiting its action (48, 49). Release of Cr from intracellular stores would decrease the measureable concentration of Cr.

A trend towards a decrease in absolute concentration of myo-inositol was also observed in the current study. To our knowledge only one other study examined a change in myoinositol after tDCS. Contrary to our results, Rango *et al.* observed an increase in the concentration of myo-inositol 30 minutes after 1.5 mA of anodal tDCS to the right motor cortex (12). The use of different current levels, electrode montage, and area measured, could all contribute to this discrepancy.

The observed association between ΔNAA and ΔtCr indicates that individuals that had a decrease in NAA following tDCS relative to sham also had a decrease in tCr levels (Figure 4). As reported earlier, reduction in NAA upon neural activation is due to increased demand for NAAG, a modulator of GABA and glutamate. As Cr and NAAG are both released from the neuron in an action-potential dependent manner to act as neuromodulators, their correlated decrease in concentration is feasible. These data support the notion that tDCS increases cortical activation, resulting in an increased neuronal energy demand, which subsequently decreases tCr and NAA.

The after effects of tDCS are thought to be dependent on alterations of the membrane potential and changes in glutamate and GABA signalling, relating to synaptic plasticity (3, 6). As such, we expected to observe changes in glutamate and GABA following tDCS. However, the literature presents conflicting results (7, 10–12, 17, 19, 50, 51). In a study observing metabolite concentration both during and after tDCS, Bachtiar *et. al.* observed a significant decrease of GABA concentration in left M1 after anodal tDCS to the same area compared to sham, but no differences between sham and anodal tDCS *during* the stimulation period (50). This indicates that the alteration of neurotransmitters that occurs due to tDCS is predominantly evident after the stimulation period. It is possible that our measurement of metabolite concentration occurred outside the optimal window of neurotransmitter modulation, and instead we observed upstream events. Further studies are required to identify the mechanism by which GABA and Glu are altered to enhance or depress synaptic activity and the optimal time to observe the peak change in these neurotransmitters.

The current study is the first to measure metabolite concentrations of the motor cortex using a bihemispheric montage in an ultra-high-field MRI (7T). Currently, there are only two studies that have examined the metabolism of the motor cortex following tDCS at 7T, both using the conventional M1-supraorbital (unihemispheric) montage, and both stimulating outside of the scanner (7, 18). Both studies examined the effects of 1 mA of cathodal stimulation over left M1 for 15 (18) and 10 (7) minutes with differing results. Stagg *et. al.* found a decrease in Glu/Cr after cathodal stimulation, while Kim and colleagues reported no significant change in Glu concentration following cathodal stimulation. Kim *et. al.* did report a significant reduction in GABA following anodal stimulation, and no changes in other key metabolites (18). The current study applied 2 mA of current for 20 minutes. It has recently been shown that cathodal stimulation, which is believed to be inhibitory, reverses its polarity at 2 mA and becomes excitatory (9). The higher current used in our study compared to previous studies may explain the differing results. Increasing the current to 2 mA delays the time of peak metabolic change from immediately after stimulation, to up to 90-120 minutes after stimulation (9). This delay may explain why we did not observe changes in Glu and GABA in the current study.

Only one other study has examined bihemispheric (M1-M1) tDCS on motor cortex metabolism (19). Using this montage, and 1mA of current for 20 minutes, Tremblay and colleagues reported no significant modulation in any metabolite concentration at 3T (19), consistent with the current study. They concluded that the complex relationship between excitatory and inhibitory mechanisms within and between the primary motor cortices resulted in high inter individual variability and response to tDCS stimulation. The ultra-high field MRS used in the current study provided greater signal to noise ratio and spectral dispersion compared to Tremblay *et al.,* (19) increasing metabolite measurement precision. This greater measurement precision may be responsible for the observation of trends towards increased NAA/tCr and decreased tCr in the current study.

Although there have been few studies observing the metabolic and functional changes following M1-M1 tDCS, the current study is the first to incorporate the SMA as a potential target for bihemispheric tDCS. The SMA has been studied as a potential tDCS target in behavioural studies of posture and visuomotor learning (28, 52). Its anatomical positioning and strong connections to M1 make it a well-suited target for motor network modulation. In addition, the SMA has strong efferent connections to the corticospinal tract, making it an ideal candidate as a target for neurorehabilitation (27). Various studies have shown the importance of the SMA and associated non primary motor areas after brain or spinal cord injury (24–26). Neural recruitment is an important aspect of recovery. SMA should be considered to enhance synaptic connections of bilateral M1, subcortical structures, and further downstream to the corticospinal tract.

One important limitation of the current study was the omission of a within session baseline measurement. The MRS data acquired for this study was part of a longer imaging protocol that incorporated anatomical and resting-state fMRI measurements (to be published elsewhere). Therefore, time constraints prevented the inclusion of a baseline spectroscopy measurement. However, the cross-over design of this study including a separate spectroscopy measurement during sham stimulation on a separate day has previously been shown to be an acceptable approach (30). However, the inclusion of a baseline measurement in the future would likely reduce inter-subject variability. A recent study at 7T has provided estimates of the reliability of metabolite measurements taken on separate days (31). In addition, a comparable protocol, using a sham control without baseline measurements observed no changes in metabolite concentration after 1 mA of bilateral dorsolateral prefrontal cortex (DLPFC), measured from the left DLPFC (11). Another limitation of the current study is that metabolite measurements were not made in the SMA, again due to time constraints. Future studies would also benefit from examining metabolite changes in this brain region following stimulation. Finally, the current study was not designed to examine GABA levels. Although, previous studies have shown changes in GABA concentration, we were not able to measure GABA with sufficient reproducibility using the spectroscopy method applied in the current study. Future studies using GABA editing methods could help elucidate the modulation of GABA by tDCS.

In conclusion, bihemispheric transcranial direct current stimulation with anode over SMA and cathode over M1 was safely applied during 7T MRI for 20 minutes at 2 mA. Immediately following stimulation there were no changes in metabolite levels measured by ^1^H MR spectroscopy of the left primary motor cortex in this sham controlled cross-over study.

However, when comparing stimulation to sham conditions, there was a significant positive association between the change in *N*-acetyl aspartate and the change in creatine in the same region.

